# Atomic Layering Thermostable Antigen and Adjuvant (ALTA^®^) platform provides unique antigen delivery system through controlled release to improve immune response to vaccination

**DOI:** 10.64898/2026.01.05.697591

**Authors:** Keith A. Strand, Heather D’Angelo, Ashley M. Gerwing, Isabella R. Walters, Emma M. Snyder, Alyssa M. Ritter, Emily Hite, Yalini H. Wijesundara, Annie B. Caplan, Daria L. Ivanova, Isabella Allen, Yu Han, Lorena R. Antunez, Antu K. Dey, Bryan L. Steadman, Sky W. Brubaker

**Affiliations:** VitriVax, Inc., 5500 Central Avenue, Boulder, Colorado, 80301, USA

**Keywords:** Vaccines, microparticles, ALTA^®^, sustained release, thermostability, antibody response, N332-GT5 gp140, HIV-1 vaccine

## Abstract

Prophylactic vaccines are commonly delivered using a multi-dose regimen with the goal of generating potent, durable protection against a specific pathogen. However, the requirement for multiple administrations can impede patient adherence and reduce overall protection. Designing a single-shot vaccine without compromising efficacy could significantly improve vaccine adherence and performance. Previously, it has been shown that atomic-layer deposition (ALD) of alumina (Al_2_O_3_) can be applied to spray dried, thermostabilized antigen-containing powders to produce alumina-coated vaccine particles that, when compared to a liquid control, elicit improved humoral immunity with response kinetics controlled by ALD-coat thickness. However, previous studies have not defined the particle release/antigen delivery profile of ALD-coated vaccines. The studies in this manuscript were designed to investigate how the kinetics of antigen release from ALD-coated vaccines impacts the timing and magnitude of the immune response relative to single- and multi-dose liquid vaccine regimens using two distinct antigens, Ovalbumin and the HIV-1 envelope trimer, N332-GT5 gp140. By combining longitudinal *in vivo* imaging and immunological readouts, we demonstrate that ALD-coated vaccines exhibit tunable, variable-rate release and deliver antigen in a unique, prolonged manner that results in an improved immune response to single-shot vaccination for difficult to target pathogens, such as HIV-1. Furthermore, using *in vitro* analytical methods, we confirmed the ability of our Atomic Layering and Thermostable Antigen and Adjuvant (ALTA^®^) platform to impart thermostability upon the N332-GT5 gp140 antigen, a clinically relevant HIV-1 immunogen, indicating the potential for ALTA^®^ formulation to generate thermostable, single-dose vaccine products.

**Highlights:**

– ALTA® microparticles provide sustained antigen delivery with variable release rates, which can be controlled by altering ALD-coat thickness
– Sustained antigen release from thermostable, spray-dried ALTA® vaccine products impacts kinetics of humoral and cellular responses, and improves antigen-specific immunogenicity compared to single administration of liquid vaccine
– ALTA® formulation imparts vaccine thermostability through spray-drying and ALD-coating to clinically relevant HIV-1 Env antigen, N332-GT5 gp140

## Introduction

Atomic layer deposition (ALD) technology is a method to apply metal oxides in atom-thick layers on a surface that is widely used to manufacture materials across various industries, but most prominently in the production of semiconductors. Spray drying techniques, typically used in the production of pharmaceuticals or the food industry, have recently been applied to vaccines to create spray dried powders containing vaccine components, both antigen and adjuvants. Our Atomic Layering Thermostable Antigen and Adjuvant (ALTA^®^) technology platform combines those two technologies to generate spray dried vaccine powders that are subsequently coated with alumina (Al_2_O_3_) using ALD. Each cycle of ALD imparts a 2.3-Å-thick layer of alumina on the spray dried powder, and by controlling the number of ALD cycles applied, we can successfully and consistently generate ALTA^®^ vaccine products with different alumina coat thicknesses[1,2].

An important feature of spray dried, ALD-coated vaccine powders is the thermostability imparted upon the antigen payload due to the spray drying process. Previously, antigen thermostability within both uncoated and ALD-coated spray dried vaccine products has been shown by demonstrating that they retain *in vivo* immunogenicity following exposure to elevated thermal conditions[3,4]. However, to our knowledge, there have been few reports directly monitoring antigen stability after ALD coating. Here, we demonstrate the ability to recover antigen from ALD-coated materials and confirm antigen integrity using both the model antigen, ovalbumin (OVA) and a clinically relevant Human Immunodeficiency Virus type 1 (HIV-1) vaccine immunogen, N332-GT5 gp140[5,6].

Previous studies have shown the potential for ALD-coated vaccines to elicit strong immune responses wherein the kinetics of the immune response are controlled in a tunable manner, with increased ALD-coat thickness correlating with delayed antibody responses *in vivo*[1–4]. Due to the observed delay in humoral response kinetics, ALD coated vaccines appeared to release their payload in a pulsatile fashion at delayed timepoints, with the timing of release being controlled by ALD coat thickness. Hence, materials of different ALD-coat thicknesses might be mixed with the intention to provide multiple vaccine doses within a single administration, wherein low ALD-coated particles release antigen earlier while the higher ALD-coated particles release antigen in a delayed manner. To that end, a single dose of ALD-coated vaccine powder, containing a mixture of particles with multiple coat thicknesses, was recently reported to elicit a more broadly neutralizing antibody (bnAb) response against SARS-like betacoronaviruses than a multi-dose liquid vaccine regime containing the same Spike antigen[7]. However, in preliminary *in vivo* imaging studies, vaccine particles coated with 250 ALD cycles persisted at the site of injection for up to 14 weeks, raising the possibility that antigen could be released over that entire duration[2]. The studies described herein were performed to provide more insight into the pharmacokinetics of antigen release from ALD-coated vaccine particles of different coat-thicknesses, with the goal of enabling ALTA^®^ vaccine product development for clinical evaluation.

The ability to manufacture and develop single-shot, thermostable vaccine products, without compromising efficacy compared to a multi-dose regimen, could provide significant public health benefits, especially in low- and middle-income countries where individuals may have less access to medical care and adherence to multi-dose vaccine regimens for full vaccine efficacy has been a challenge. Recently, the germline-targeting antigen N332-GT5 gp140 and a potent saponin/monophosphoryl lipid A nanoparticle (SMNP) adjuvant have been shown to prime precursor B cells with the potential to evolve into bnAb producers against HIV-1 in non-human primates (NHPs) using a seven-administration, escalating dose vaccination paradigm that is now being evaluated in a human clinical trial[6,8,9]. Despite the encouraging results from this dosing regimen in NHPs, the need to administer seven priming injections of N332-GT5 gp140 + SMNP over the course of 12 days makes the regimen practically challenging. Given the recent demonstration that single-dose ALD-coated vaccine particles can elicit a robust bnAb response against sarbecoviruses[7], this HIV-1 antigen-adjuvant regimen, requiring multiple priming doses and subsequent boosts, may particularly benefit from formulation within the ALTA^®^ vaccine platform to limit the number of administrations required.

Here, by combining longitudinal *in vivo* imaging, immunological readouts, and analytical test methods, we provide insight into how the ALTA^®^ platform may be leveraged to generate vaccine products possessing unique antigen delivery kinetics that elicit a robust immune response following single-shot vaccination.

## Methods

### Manufacture of ALTA^®^ materials

Spray-dried powders were made using a Buchi Mini B-290 spray dryer with a B-296 Dehumidifier (BUCHI, New Castle, DE, USA). Spray-drying setpoints were chosen to manufacture powders with the desired size and moisture content characteristics while maximizing yield. The proprietary liquid formulations utilized for spray drying contained 17% w/v total solids, including ovalbumin (OVA) (InvivoGen #vac-pova-100) or N332-GT5 gp140 (supplied by International AIDS Vaccine Initiative (IAVI)) at 1% w/w (solids basis), or 2-[(1*E*,3*E*)-4-[4-(Dimethylamino)phenyl]-1,3-butadien-1-yl]-4,5-dihydro-(4*S*)-4-thiazolecarboxylic acid hydrochloride salt (AkaLumine-HCl, TokeOni) (Sigma Aldrich #808350 or MedChemExpress, #HY-112641A) at 2.0-2.5% w/w (solids basis). The intermediate spray-dried powder was coated with aluminum oxide in a mechanically agitated fluidized-bed ALD reactor, using trimethylaluminum (Strem #93-1360) and water as precursors[2,3]. Purge steps were employed after each precursor exposure to avoid chemical vapor deposition. The coating was performed at 50 °C, using nitrogen as the fluidization gas. The number of ALD cycles used to coat powders in these studies were 50, 100 or 200.

### Fluorescent labeling of OVA and N332-GT5

Stock solutions were prepared of ovalbumin (InvivoGen #Vac-stova Lot #5823-45-01) and N332-GT5 gp140 (IAVI) at approximately 10 mg/mL in 50 mM sodium borate, pH 8, and IVISense680-NHS (Revvity #NEV11120) at 18-27 mM in DMSO. The IVISense680-NHS stock solution was spiked into the antigen stock solutions at a dye:antigen molar ratio of 8.85:1 (OVA) or 10:1 (N332-GT5). The mixture was protected from light and incubated at room temperature for 2-4 hrs then moved to 4 °C overnight. Following incubation, excess free IVISense680-NHS dye was removed using a 7k MWCO Zeba Desalting Column (Thermo Scientific #89893) per manufacturer instructions and conjugated antigens were further purified and buffer exchanged by dialysis.

### Mechanical disruption of ALTA^®^ powders

To acquire broken ALTA^®^ particles, intact ALTA^®^ powder was mechanically disrupted to break the alumina shell. To confirm OVA remained intact after mechanical disruption, broken ALTA^®^ powder and spray dried intermediate (precursor to ALTA^®^ OVA powder) were reconstituted in PBS/0.05% Tween-20 at 10 mg/mL. Size Exclusion Chromatography (SEC) was performed on the Waters ACQUITY UPLC Instrument. Mobile phase: isocratic flow of 100 mM Sodium Phosphate (pH 6.80 ± 0.05) at 0.30 mL/min, column: GTX450 SEC Column (Waters), injection volume: 10 µL.

### N332-GT5 trimer Potency by ELISA

ALTA^®^ particles were mechanically disrupted, and contents were reconstituted in PBS/0.05% Tween-20 at 10 mg/mL for analysis by an indirect, enzyme-linked immunosorbent assay (ELISA) to determine anti-N332-GT5 potency. Nunc 96-well flat bottom high-binding plates (Thermo Scientific #442404) were coated with 0.1 µg lectin (Sigma #L8275) in PBS overnight at 4 °C, then incubated with a blocking buffer (2% BSA in PBS) at room temperature for 2 hrs. Then, plates were coated with 0.2 µg of recovered sample or recombinant N332-GT5 gp140 trimer reference material (supplied by IAVI) and incubated for 1 hr at 37 °C. A standard curve was established using a dilution series of BG18 bnAb (supplied by Scripps Research Irvine & Stiechman Labs). The reference material or test samples were incubated for 1 hr at room temperature, followed by a wash step. Horseradish peroxide-conjugated goat-anti-human secondary antibodies (BioRad #5172-2504) were added at the manufacturer-specified dilution and incubated for 1 hr at room temperature. Excess secondary antibody was washed off, and the plates were developed for 5 minutes using Ultra TMB (Thermo Fisher #34028) and quenched using sulfuric acid (H_2_SO_4_). Absorbance was measured at 450 nm and 650 nm on a BioTek Synergy plate reader (Agilent) within five minutes after the addition of H_2_SO_4_. To analyze the data, 650 nm absorbance was subtracted from the 450 nm absorbance and plotted against the log(10) of primary antibody concentration. The potency for each sample was determined by comparing the total area under the curve (AUC) for each sample to the reference material AUC from each plate using GraphPad Prism (version 10.4.0).

### *In vivo* vaccine administration and serum collection

Vaccine Administration and Animal Studies were conducted under approval from the Institutional Animal Care and Use Committee at the University of Colorado, Boulder, CO (protocol #2835 and #2838). Mice were prepped for vaccination by shaving the right hindlimb to remove fur. ALTA^®^ powders were resuspended at the indicated concentrations in sterile 6% hetastarch + 0.9% saline (Hospira NDC: 00409-7248-03) or Super Refined™ Sesame Oil (Croda Cat. #SR40294) prior to injection. Alhydrogel^®^ (InvivoGen #vac-alu-50) and OVA (InvivoGen #Vac-stova Lot #5823-45-01) used in the animal studies were prepared following the manufacturer’s instruction. Briefly, Alhydrogel^®^ adjuvant 2% was combined with the OVA antigen in a 1:1 volume, mixed, and allowed to adsorb for 30 min at room temperature prior to administration. Monophosphoryl Lipid A (MPLA) + Alhydrogel^®^ diluent was made by combining MPLA-SM (InvivoGen #tlrl-mpla) and Alhydrogel^®^ at a 1:10 ratio and mixing by rotation for 1 hr at room temperature. The Alhydrogel^®^-adsorbed MPLA was pelleted by centrifugation, then resuspended in sterile 6% hetastarch + 0.9% saline. This mixture was used to resuspend ALTA^®^ particles prior to administration. To generate liquid formulations of N332-GT5 gp140 (IAVI) + SMNP (supplied by Scripps Research Irvine Lab), stock solutions of the two components were mixed at the desired ratio and diluted with sterile 1x PBS prior to administration. SMNP diluent for ALTA^®^ products was created by diluting SMNP stock in sterile 6% hetastarch + 0.9% saline (Hospira NDC: 00409-7248-03) and used to resuspend ALTA^®^ particles. Vaccine administrations were given via a single 50 µL intramuscular injection to the right hindlimb. Immunogenicity studies using the antigens OVA or N332-GT5 gp140 were performed using 6-8 week old, female C57BL/6J mice (Jackson Labs, strain #000664). Longitudinal fluorescent imaging studies were performed using 6-8 week old, female SKH-1 Elite mice (Charles River Laboratories, strain #477), while longitudinal bioluminescent imaging studies were performed using male and female FVB-Tg(CAG-luc,-GFP)L2G85Chco/J (Jackson Labs, strain #008450) [18] that were bred internally and aged between 6-12 weeks at the time of ALTA^®^ administration. Blood was collected via submandibular bleed and serum isolated using gel separation tubes (Sarstedt, Nümbrecht, Germany, #41.1378.005) and stored at –80 °C prior to analysis.

### Serum antibody titer measurement by ELISA

#### Anti-OVA IgG1

Mouse serum samples were analyzed by an ELISA to determine anti-OVA IgG1 antibody titers. Nunc 96-well flat bottom high-binding plates (Thermo Scientific #442404) were coated with 5 µg/well of OVA (Fisher #BP2535-5) in phosphate-buffered saline (PBS), incubated at 4 °C overnight, and then rinsed four times using a wash buffer (0.05% Tween 20 in PBS). The plates were then incubated with an assay-blocking buffer (3% BSA, 0.05% Tween 20 in PBS) at room temperature for one hour. A standard curve was established with a dilution series of mouse anti-OVA IgG1 primary antibody (Chondrex #7093). The calibrated primary antibody and mouse sera samples were incubated for 1 hour at room temperature, followed by a wash step to remove the unbound antibody. Horseradish peroxide-conjugated secondary antibody (Southern Biotech #1071-05) was added at 1:4000 dilution and incubated for one hour at room temperature. Excess secondary antibody was washed off, and the plates were developed for 20 min using Ultra TMB (Thermo Fisher #34028) and quenched using H_2_SO_4_. Absorbance was measured at 450 nm and 650 nm on a BioTek Synergy plate reader (Agilent) within five minutes after the addition of H_2_SO_4_. The standard curves and interpolated data for each plate were independently generated using Gen5 software (Agilent).

#### Anti-N332-GT5 gp140 total IgG, IgG1, IgG2a, IgG2b, and IgG3

Mouse serum samples were analyzed by an indirect ELISA to determine anti-N332-GT5 gp140 antibody titers. Nunc 96-well flat bottom high-binding plates (Thermo Scientific #442404) were coated with 2 µg/mL lectin (Sigma #L8275) in PBS for 4 hrs at room temperature, then incubated with an assay-blocking buffer (2% BSA in PBS) at 4 °C overnight. Then, plates were coated with 50 µg recombinant N332-GT5 gp140 trimer (supplied by IAVI) and incubated for 1 hr at room temperature. Excess trimer was removed with a wash step, as described above. A standard curve was established using a dilution series of positive mouse sera (supplied by Scripps Research Irvine Lab). The positive serum controls or test samples were incubated for 1 hour at room temperature, followed by a wash step. Horseradish peroxide-conjugated goat-anti-mouse secondary antibodies (**total IgG 1:5000:** BioRad #1721011, **IgG1 1:12000**: BioRad #STAR132 **IgG2b 1:12000** BioRad #STAR134 **IgG2c 1:2000** BioRad #STAR135 **IgG3 1:10000** BioRad #STAR136) were added at the specified dilutions and incubated for 1 hr at room temperature. Excess secondary antibody was washed off, and the plates were developed for 20 min using Ultra TMB (Thermo Fisher #34028) and quenched using H_2_SO_4_. Absorbance was measured at 450 on a BioTek Synergy plate reader (Agilent) immediately after the addition of H_2_SO_4_. For the standard curve, the first dilution of positive control sera was assigned an arbitrary value of 10,000 units and subsequent dilutions of positive control sera were given corresponding values based on their dilution factor. The interpolated data from each plate was independently generated using Gen5 software (Agilent).

### *In vivo* Fluorescent Imaging and Analysis

Fluorescent images were acquired using the IVIS Lumina III imaging system with Living Image software (version 4.8.0, Revvity) housed in the JSCBB vivarium at the University of Colorado-Boulder. Mice were anesthetized using inhaled isoflurane. Fluorescent imaging throughout the study was performed using 660/710 nm excitation/emission filters for VivoTag-680 dye. Image settings for all time points were kept constant as follows, FOV: E with XFOV-24 lens, Exposure: Auto, Pixel Binning:8, F/Stop: 2. Imaging was performed immediately post injection, and at 6 and 24 hrs post-injection. Following this, mice were imaged regularly throughout the remainder of the studies. Image data was analyzed using the Living Image software to quantify radiant efficiency [p/sec/cm^2^/sr]/[µW/cm^2^] within a fixed region of interest (ROI) at all timepoints. To quantify signal loss over time, radiant efficiency at each timepoint was expressed as a percentage relative to the radiant efficiency measured within the ROI at 24 hrs post injection. Non-linear (4PL) regression analysis of the data was performed using GraphPad Prism (version 10.4.0).

### *In vivo* Bioluminescent Imaging and Analysis

Images were acquired using the IVIS Lumina III imaging system with Living Image software (version 4.8.0) housed in the JSCBB vivarium at the University of Colorado-Boulder. Mice were anesthetized and imaged regularly as described above. Imaging settings were kept constant throughout all studies, with settings as follows: excitation/emission: block/open, FOV:E with XFOV-24 lens, exposure: 60 sec, pixel binning:8, F/Stop: 1.2. Radiance (p/sec/cm^2^/sr) was measured within a fixed ROI using Living Image software. To quantify luminescence from D1 to study end, the total area under the curve (AUC) of luminescent signal from each animal was calculated using a trapezoidal method. The percentage of AUC measured per day was calculated as follows: Percent AUC = (daily AUC)/(total AUC)*100. This data was then summed and analyzed using a non-linear (5PL) regression performed in GraphPad Prism (version 10.4.0). The first derivative of the 5PL curve was generated using GraphPad Prism.

### Flow cytometry for identification of OVA-specific CD8+ T cells in blood

Whole blood was collected into the EDTA-treated tubes (EDTA K3E, Sarstedt). The blood samples were RBC lysed (ACK Lysing Buffer, Thermo Scientific) for 5 min at RT, quenched, and washed with complete media (RPMI1640 with L-glutamate supplemented with 10% fetal bovine serum, penicillin/streptomycin, sodium pyruvate, non-essential amino acids, beta-mercaptoethanol, 4-(2-hydroxyethyl)-1-piperazineethanesulfonic acid (HEPES)). Cell suspensions were plated onto the U-bottom 96-well plate and incubated with αCD16/32 to block Fc receptors. To stain OVA-specific CD8+ T cells, cell suspensions were incubated with CD8a (53-6.7, BioLegend) and tetramers (Flex-T™ Biotin H-2 K(b) OVA Monomer (SIINFEKL) (BioLegend) tetramerized by Streptavidin-BV421™ or Streptavidin-APC (BioLegend) according to the manufacturer’s protocol) for 30 minutes at 37 °C. Cells were then washed with complete media and stained with the viability dye (Ghost Dye Red 780 Fixable Viability Dye, Cell Signaling Technology), CD19 (6D5, BioLegend), CD44 (IM-7, Tonbo) in Flow Staining Buffer (Cytek) for 20 minutes RT. After washing with Flow Staining Buffer, the samples were fixed for 30 minutes in Fixation Buffer (Cytek), washed again, and resuspended in Flow Staining Buffer. Samples were acquired on Cytek Northern Lights 3-laser (VBR) spectral flow cytometer and analyzed using FlowJo 10.10.0 software (BD Biosciences), Excel (Microsoft), and GraphPad Prism (version 10.4.2).

## Results

### Kinetics of humoral response are controlled by ALD-cycle number and dependent upon dose administered

Previously, using powders coated with 50-1000 ALD cycles, we demonstrated that by increasing the number of ALD cycles we could generate delayed IgG1 responses at a specific dose of the model antigen OVA[1]. The observed delay in time to seroconversion (defined as 2 log-fold increase in anti-OVA IgG1 titer relative to pre-injection baseline) correlated with the *in vitro* dissolution assay results, suggesting the possibility of achieving controlled, delayed antigen release without significant loss in immunogenicity from powders coated with 250-500 ALD cycles[1]. While encouraging, these findings were based on a single antigen dose (250 ng OVA) achieved by administering a single 50 µL injection of ALD-coated powders at approximately 0.5 mg/mL concentration. Antigen percentage can be controlled within our formulation to create powders with different properties, however, the percentage of antigen within a specific ALTA^®^ product is constant; therefore, to achieve increased antigen doses with the same powders at a constant dose volume, we must administer ALTA^®^ powder at higher mg/mL concentrations. Here, we were interested in testing whether we would observe a similarly delayed immune response from 250+ ALD-cycle material at higher antigen doses by simply increasing the amount of ALD-coated powder administered. The *in vitro* dissolution assay results confirmed that increasing ALD-cycle number increased time to inflection (50% release) for these powders, suggesting delayed antigen release (Supplementary Table 1). Similar to previous results, we found that 50-cycle material elicited >50% anti-OVA IgG1 seroconversion by week 2 at all doses tested, and that at equivalent doses, a delay in time to >50% seroconversion was observed for 250-cycle and 400-cycle powder (Supplementary Figure 1A-1C). However, we observed that the time to reach >50% seroconversion for 250 and 400 ALD-cycle material was dose dependent, where higher antigen doses (higher mg/mL powder concentrations) elicited seroconversion at earlier timepoints (Supplementary Figure 1A-1C). When we correlated the *in vitro* time to 50% release for each powder with the time to seroconversion for individual animals at each dose, we found that the slope of the regression line was lower at the 1000 ng dose than at the 333 ng or 111 ng doses, indicating that the observed delay in time to seroconversion is reduced at higher antigen doses (Supplementary Figure 1D). These results demonstrate that increasing the amount of powder administered can accelerate the onset of the immune response regardless of the number of ALD-cycles.

**Figure 1.**
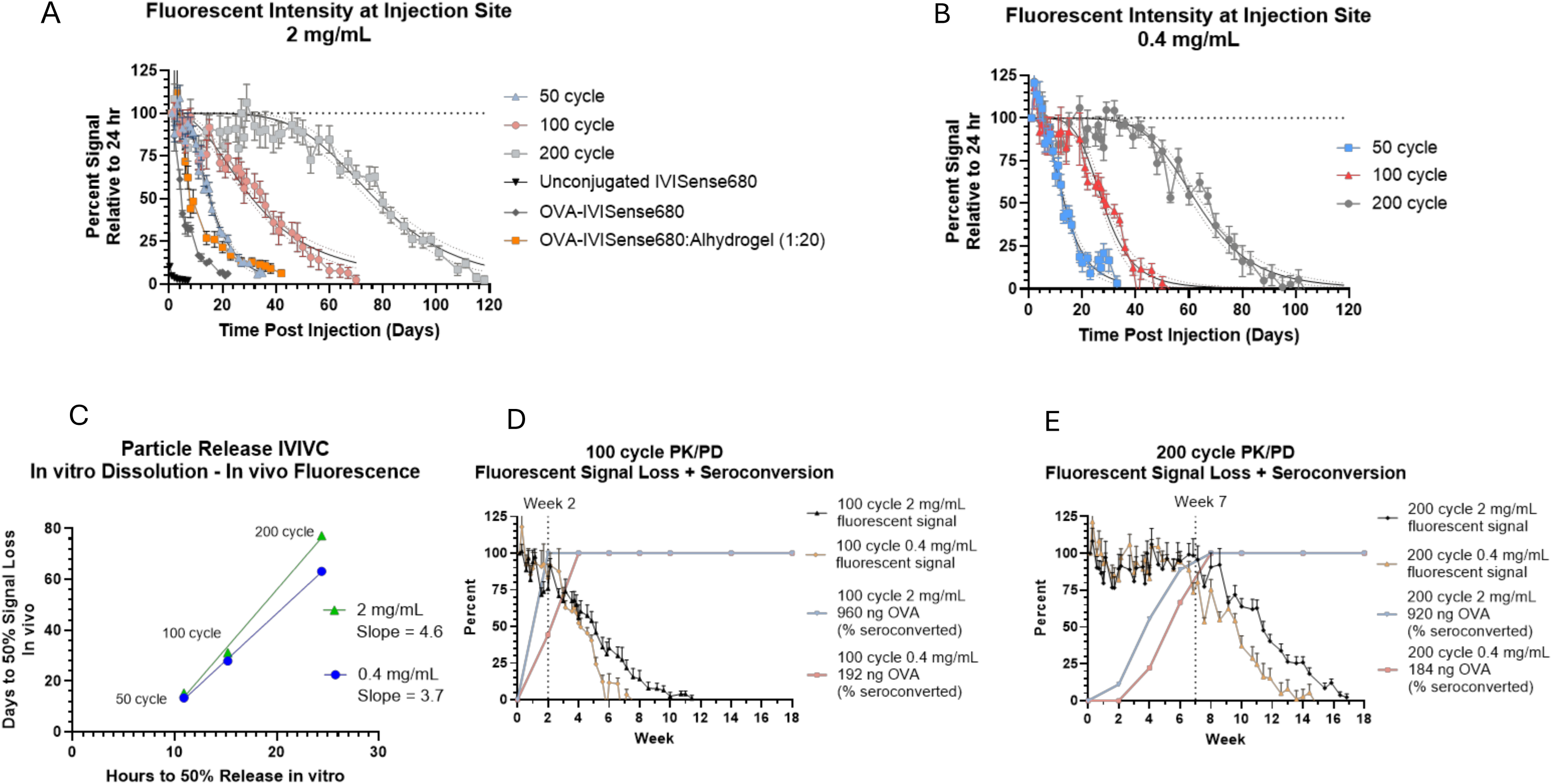
– ALD coated particles persist at site of injection with duration controlled by increasing ALD coat thickness and dose administered. A-B. Group mean +/- SEM percent of fluorescent radiant efficiency (p/s)/(µW/cm^2^) relative to radiant efficiency at 24 hr timepoint measured at site of injection for 50-, 100- or 200-cycle ALD coated powders administered at 2 mg/mL or (B) 0.4 mg/mL. Unconjugated, liquid IVISense680 fluorescent dye, conjugated OVA-IVISense680 or Alhydrogel-adsorbed, conjugated OVA-IVISense680 were administered at equimolar concentrations to amount of fluorescent dye in ALD coated powder at 2 mg/mL dose. Non-linear 4PL regression fit line with 95% confidence intervals shown for ALD coated powders (constraints on 4PL regression bottom = 0, top = 100) C. Correlation of time to 50% particle dissolution in vitro and time to 50% fluorescent signal loss in vivo (using 4PL fit parameters) at indicated doses. Simple linear regression of data at each dose shown, with slope of regression line indicated on plot. D-E. Anti-OVA IgG1 seroconversion percentage following vaccination with 100-cycle ALTA^®^ OVA dosed at 2 mg/mL (960 ng OVA) or 0.4 mg/mL (192 ng OVA) or (E) 200-cycle ALTA^®^ OVA dosed at 2 mg/mL (920 ng OVA) or 0.4 mg/mL (192 ng OVA). The IgG1 data is overlaid with group mean +/- SEM percent of fluorescent radiant efficiency relative to 24 hr timepoint measured at site of injection. Dotted lines at week 2 or week 7 indicate shift in rate of fluorescent signal loss for (D) 100-cycle or (E) 200-cycle powders, respectively.

### *In vivo* imaging demonstrates sustained, variable-rate release from ALD-coated powders

To further understand the observed dose-dependent pharmacodynamics, we generated 50-cycle, 100-cycle and 200-cycle ALTA^®^ particles containing OVA labeled with IVISense680 fluorescent dye which were used to assess *in vivo* particle release kinetics by performing longitudinal *in vivo* imaging. Using this method, we were able to quantify fluorescent signal loss at the injection site over time, which we interpret as particle release or antigen delivery. The 50-, 100- and 200-cycle ALD materials given at 2 mg/mL (0.1 mg powder mass) persisted at the injection site for 35, 70 and 118 days, respectively (Figure 1A). At 0.4 mg/mL ALTA^®^ concentration (0.02 mg powder mass), time to reach 0% signal was reduced by 2 days for 50-cycle ALTA^®^, 20 days for 100-cycle ALTA^®^, and 22 days for 200-cycle ALTA^®^ relative to a 2 mg/mL dose concentration (Figure 1B). Conversely, increasing the dose of 50-cycle ALTA^®^ to 10 mg/mL (0.5 mg powder mass) prolonged fluorescent signal at the site of injection by 22 days relative to a 2 mg/mL dose (Supplementary Figure 2A). While retention time at the site of injection was impacted by changes to the concentration administered, it was not affected by choice of diluent. Fluorescent signal loss at the injection site from 50-, 100- and 200-cycle materials dosed at 2 mg/mL was similar when administered in a non-aqueous diluent (super-refined sesame oil), indicating that the choice of diluent does not cause significant alterations to the antigen release kinetics of ALTA^®^ vaccine powders (Supplementary Figure 2B).

The persistence of fluorescently labeled ALTA^®^ particles at the injection site was much longer than liquid controls containing either unconjugated fluorescent dye or OVA-conjugated dye, at equimolar dye concentrations, which were no longer detectable at the injection site after 1 day or 21 days, respectively (Figure 1A). Interestingly, Alhydrogel-adsorbed OVA-IVISense680, which is widely believed to form an antigen depot at the injection site and is used widely as an adjuvant in clinical vaccine formulations, persists at the injection site for a similar amount of time as 50-cycle material when delivering equivalent antigen amounts despite large differences in total Al^3+^ administered per dose (19.2 µg Al^3+^ in Alhydrogel dose vs. 1.5 µg Al^3+^ in 2 mg/mL dose 50-cycle ALTA^®^ material) (Figure 1A).

To better understand the impact of dosing concentration on *in vivo* antigen release kinetics, correlations were drawn against particle release from the *in vitro* dissolution assay. A non-linear regression (4PL) model was used to analyze the fluorescence data to model time to 50% signal loss *in vivo* (4PL parameters shown in Supplementary Table 2). We then correlated the *in vitro* time to 50% release for each product (Supplementary Table 3) with the time to 50% fluorescent signal loss *in vivo* at both 2 mg/mL and 0.4 mg/mL doses. Using this correlation to predict time to *in vivo* release, a difference of 1 hour on the *in vitro* assay amounts to an additional 3.7 days in time to 50% signal loss *in vivo* at a 0.4 mg/mL dose but adds an additional 4.6 days in time to 50% signal loss *in vivo* at a 2 mg/mL dose (Figure 1C). This demonstrates that while the duration of antigen release can be tuned by controlling the number of ALD cycles applied to a given product, administering higher amounts of powder per injection can also extend the duration of particle release/antigen delivery *in vivo*.

Notably, we observed bi-phasic loss of fluorescent signal over time. At early timepoints there was a low, but non-zero, rate of signal loss, followed by a period of signal loss at a higher rate. This rate shift was observed for all ALD-coated products but was most noticeable with 200 coat material (Figure 1A-1B). The duration of the lower release rate window increased with increasing ALD-cycle number, as a shift to higher release rate was observed at D4-D8 with 50-cycle material, D14 with 100-cycle material, and D49 with 200-cycle material (Figure 1A). Similarly, the period of maximal release was extended by increasing ALD-coat thickness, as peak antigen release from 50-cycle material lasted for ∼28 days (D5-D33) but persisted for ∼70 days (D50-D118) with 200-cycle material (Figure 1A). When comparing fluorescent signal loss (antigen release) with time to seroconversion, it is evident that some antigen is delivered during the early period of lower release rate, given that at the 2 mg/mL dose, we observed >80% seroconversion in anti-OVA IgG1 titers by week 2 for 100-cycle material and week 6 for 200-cycle material while the fluorescent signal remained above 80% in both cases, indicating that the majority of antigen had yet to be released (Figure 1D-1E). Again, we observed a dose-dependent delay in time to anti-OVA IgG1 seroconversion, with the 0.4 mg/mL dose eliciting >80% seroconversion 2 weeks later than the 2 mg/mL dose, which correlated with increased fluorescent signal loss (antigen release) during this time (Figure 1D-1E).

One caveat to assessing particle release by quantifying fluorescent signal at the injection site is that signal loss may not equate directly with particle release, as particles may traffic away from the injection site to distal regions of the body undetected. In these experiments, we did not detect a significant fluorescent signal at regions other than the injection site, so while particle trafficking may occur, it appears that the majority of particles remain at the injection site. Alternatively, antigen may be released from a particle but then remain at the injection site for some time, similar to what is observed following administration of free OVA-dye conjugate (Figure 1A). To address these potential issues and generate an activatable probe to measure particle release directly, we manufactured 50-cycle, 100-cycle and 200-cycle ALTA^®^ particles containing AkaLumine-HCl (TokeOni), a D-luciferin analog with a luminescence peak in the near-infrared rage (670-680 nm) to enable deep tissue *in vivo* imaging[10], and administered these particles in a single 50 µL IM injection to FVB-Tg(Cag-luc,-GFP)L2G85Chco/J (L2G85) mice that constitutively express firefly luciferase in almost all tissues. Following administration with 100-cycle material, we observed that there was a detectable, dose-dependent luminescent signal at the injection site 20 minutes after administration, indicating the presence of some immediately soluble TokeOni following suspension of ALD-coated particles in liquid injection buffer (Supplementary Figure 3). When administered in a liquid solution at 0.4 mg/mL or at a 20 mg/mL dose of spray-dried intermediate to dose-match the amount of TokeOni in ALTA^®^ powder dosed at 20 mg/mL, TokeOni elicited a strong luminescent signal with rapid decay, displaying a ∼4 log-fold loss (0.015% of post-injection maximum) in flux by 24 hrs post injection and no detectable signal at D3 post injection (Supplementary Figure 4). Due to the observed rapid kinetics of signal decay, luminescence from ALD-coated TokeOni powder is likely generated by substrate released from ALTA^®^ particles within the prior 24 hrs.

Luminescence signal from the ALD-coated particles was detectable above pre-injection baseline levels at the injection site continuously between 20-90 days following administration of ALD-coated TokeOni powder (Figure 2A). This suggests continuous release over that time and contrasts with the transient signal observed following injection with liquid TokeOni solution or the reconstituted spray-dried TokeOni. Similar to the kinetics of ALTA^®^ containing fluorescently labelled OVA, the duration of luminescent signal was observed to be extended by both increasing ALD coat thickness and increasing particle dose (Figure 2A-2B). We then used the injection site flux data and calculated total area under curve (AUC) and percent of AUC per day to quantify particle release kinetics from 50-, 100- and 200-cycle ALTA^®^ TokeOni at 10 and 2 mg/mL (Figure 2C-2D). Using a non-linear (5PL) regression model to fit the data, we calculated the first derivative of the 5PL regression to visualize percent release per day from each product tested (Figure 2E-2F). These data clearly demonstrate how increasing ALD cycle number alters *in vivo* release kinetics, such that 50-cycle ALTA^®^ material exhibited peak release immediately upon injection, while peak releases from 100-cyle and 200-cycle ALTA^®^ materials were delayed until roughly D30 or D55 post-injection, respectively (Figure 2E-2F). Consistent with our findings using fluorescently labeled antigen, we observed variable particle release rates from 100- and 200-cycle ALTA^®^ materials, with a period of low, but non-zero, release immediately upon injection, followed by a gradual rise in release rate over time until peak release was reached (Figure 2E-2F). Together, using two independent *in vivo* imaging modalities, our data demonstrate that ALD-coated powders deliver their payload in a manner that could be described as sustained release with variable release rate.

**Figure 2.**
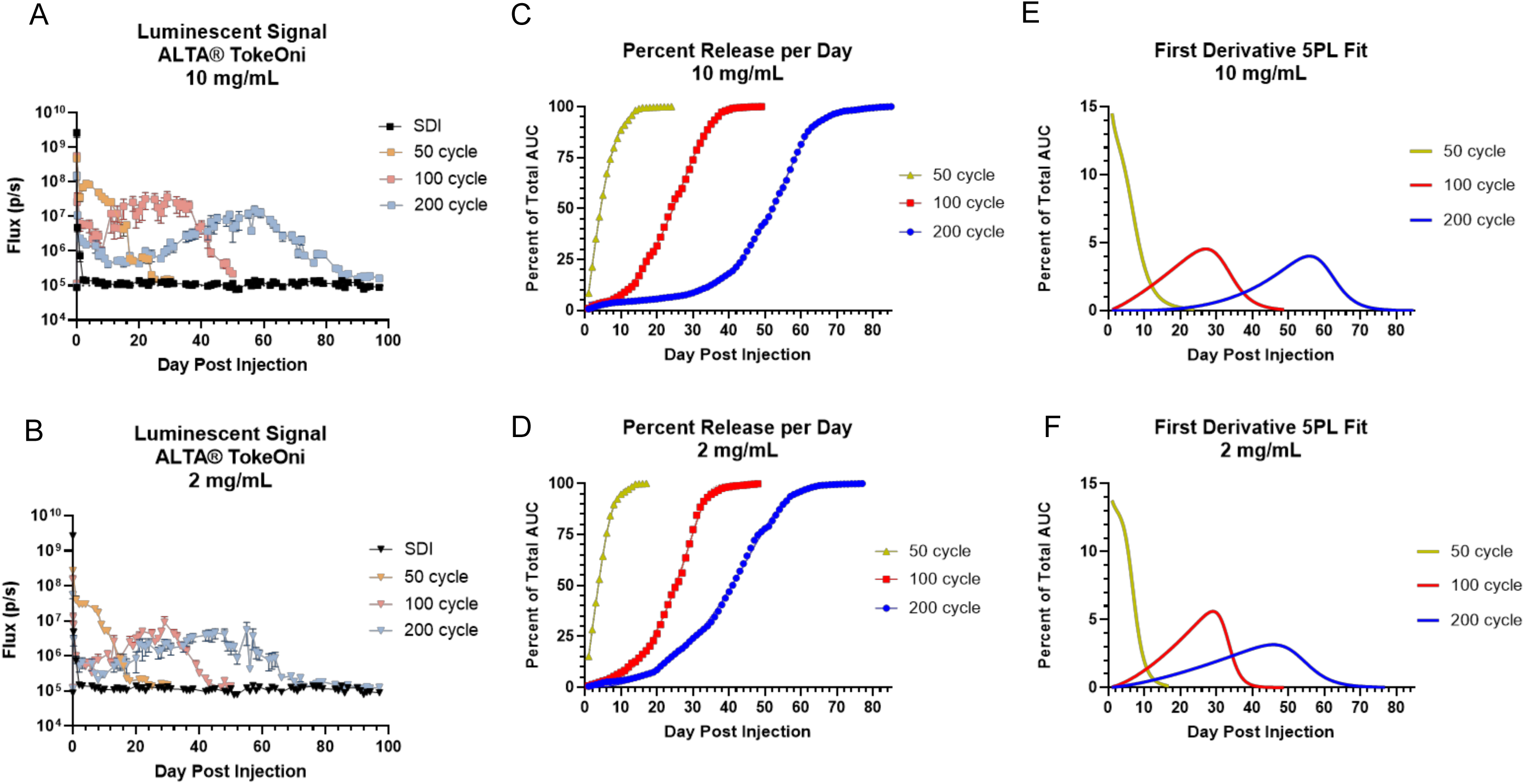
– Activatable probe demonstrates sustained particle release with variable release rate over time. A-B. Total flux (p/s) measured at the site of injection over the duration of the in vivo imaging study following administration of 20 mg/mL spray dried intermediate (SDI), and 50-, 100- or 200-cycle ALD coated TokeOni powder at 10 mg/mL dose or (B) 2 mg/mL. Dotted line represents pre-injection baseline flux at site of injection, immediately prior to injection. C-D. Using the total flux (p/s) data, the total area under the curve (AUC) was calculated for 50-, 100- and 200-cycle ALD-coated TokeOni powder at 10 mg/mL or (D) 2 mg/mL. The AUC per day was calculated, expressed as a percent of total AUC, then the daily percent AUC was summated to show cumulative percent of AUC over time. Solid line shows 5PL model fit of cumulative percent AUC data. E-F. Solid lines show the first derivative of the 5PL model fit of cumulative AUC data for 50-, 100-, and 200-cycle ALD-coated TokeOni at 10 mg/mL or (F) 2 mg/mL

### Simulating sustained antigen delivery using a multi-dose regimen of liquid vaccine partially mimics improved humoral response elicited by ALD-coated vaccines

Recently, it has been shown that sustained antigen delivery via osmotic minipump, improved antigen-alum adsorption, or altered dosing schedules can improve overall antibody production and germinal center responses[11–15]. We have previously demonstrated that ALD-coated powder elicits an improved humoral response compared to a single administration of Alhydrogel-adjuvanted liquid vaccine[1,2]. Given our results demonstrating prolonged antigen release from ALD-coated material, we hypothesized that this sustained antigen delivery profile may improve the immunogenicity of ALD-coated vaccines relative to a traditional liquid vaccine. To test this hypothesis using the model antigen OVA, with the caveat that antigen release from ALD-coated material extends beyond 7 days, we developed a rudimentary model to mimic extended antigen release by delivering a fixed OVA dose over the course of 7 consecutive days using a liquid formulation of OVA-Alhydrogel (Figure 3A), similar to previously reported extended priming strategies[12]. Additionally, to re-capitulate a liquid vaccine using our ALD-coated OVA formulation, we intentionally broke 50-cycle powder using mechanical disruption to release all encapsulated OVA prior to administration (Supplementary Figure 5). By administering this broken 50-coat material in a single shot, or over a 7-day dosing schedule with 1/7^th^ the total dose administered per day, we were able to test whether loss of antigen encapsulation to remove the sustained release properties reduces ALD-coated vaccine efficacy, and whether extended dosing of un-encapsulated antigen could rescue any potential loss in immunogenicity (Figure 3A).

**Figure 3.**
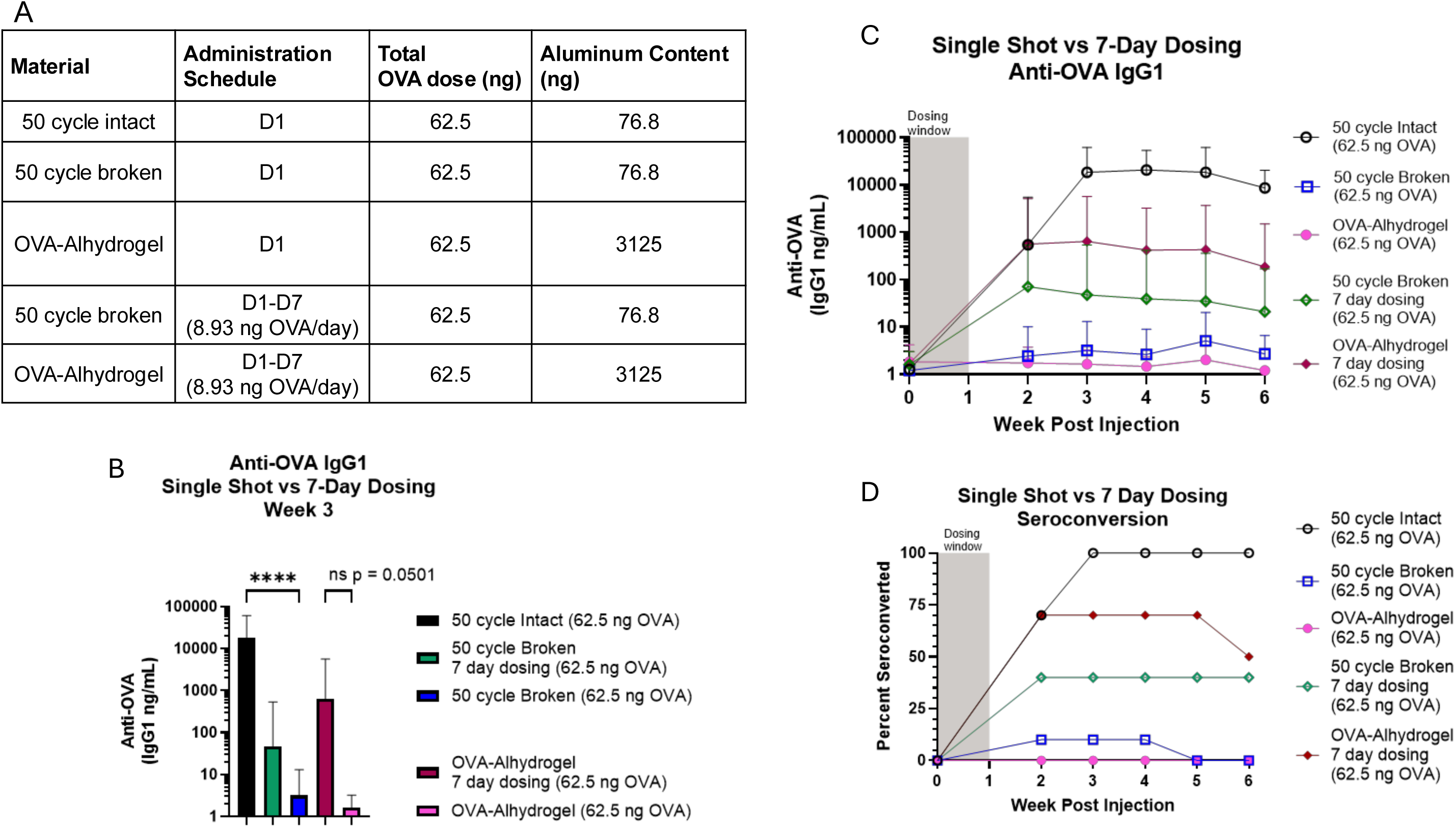
– Modeling sustained antigen delivery from ALD coated particles improves antibody response to vaccination. A. Study design, n=10 mice per group. All mice received 62.5 ng OVA dose, either in a single injection, or over the course of 7 daily injections. Of note, Al^3+^ ion content differs significantly between groups treated with ALD-coated material or Alhydrogel. B. Anti-OVA IgG1 titers at week 3 post first injection (14 days post final injection for 7 day dose groups). Plot shows geometric mean titer (n=10/group) with 95% confidence interval, **** = p < 0.0001. Data analyzed using Kruskal-Wallis test. C. Anti-OVA IgG1 titers throughout study, plot shows geometric mean titer (n=10/group) with 95% confidence interval. D. Anti-OVA IgG1 seroconversion percentage, indicating a 2 log-fold increase in anti-OVA IgG1 titers relative to pre-injection baseline.

Strikingly, while intact 50-cycle material elicited strong anti-OVA IgG1 responses and 100% seroconversion (10/10 mice) by week 3 post-injection, loss of antigen encapsulation due to mechanical disruption rendered 50-cycle material non-immunogenic with a significant reduction in anti-OVA IgG1 titers and only 10% seroconversion (1/10 mice) at this timepoint (Figure 3B-3D). As expected, intact 50-cycle material also significantly outperformed a single administration of Alhydrogel-adjuvanted liquid vaccine at a 62.5 ng OVA dose. Additionally, intact particles elicited higher anti-OVA IgG1 titers than either group given an extended dosing regimen (Figure 3B-3C). Interestingly, when compared to a single administration of liquid OVA-Alhydrogel, a 7-day dosing schedule improved anti-OVA IgG1 responses (p = 0.0501) and seroconversion percentage (0/8 mice - single dose; 7/10 mice - extended dosing) at 3 weeks post-vaccination (Figure 3B-3D). Similarly, using an extended dosing schedule improved the response at 3 weeks following vaccination with mechanically disrupted 50-cycle material, eliciting improved seroconversion rates (1/10 mice - single dose; 4/10 mice - extended dosing) and moderately increased anti-OVA IgG1 titers (Figure 3B-3D). Relative differences in the extent to which an extended dosing schedule improved the immunogenicity of OVA-Alhydrogel or mechanically disrupted ALD-coated powder may be due to the difference in total Al^3+^ administered, with the OVA-Alhydrogel groups receiving significantly more Al^3+^ than the group receiving the disrupted 50-coat ALTA^®^ material (Figure 3A). These data demonstrate that an extended vaccine dosing schedule improves humoral responses relative to a single shot, partially re-creating the strong IgG1 responses observed following vaccination with intact 50-cycle material, and hint at the possibility that improved immunogenicity of ALD-coated vaccines may be caused by their unique, sustained antigen release profile.

### Single administration ALD-coated vaccine products elicit stronger antigen specific humoral and cellular responses than two-dose liquid prime/boost vaccination

Given the prolonged, variable-rate release observed from ALD-coated vaccine powders and their ability to generate an improved humoral response relative to a single shot of liquid vaccine, we next wanted to compare the immunogenicity of ALTA^®^ vaccine powders against a traditional two-dose prime and boost vaccination schedule using the model antigen OVA. We also wanted to investigate the possibility of resuspending 100- or 200-cycle ALTA^®^ powders in a liquid vaccine formulation with the intent of providing an initial priming antigen dose from the liquid formulation, then utilizing the sustained release profile from ALTA^®^ powders to mimic a temporally delayed liquid vaccine boost. Liquid boost injections were provided on D28 or D49 post-prime, which were selected to match peak antigen release from 100-cycle or 200-cycle material, respectively (Figure 2E-2F, Supplementary Table 4).

Relative to vaccination with a two-administration liquid prime and boost, which did not elicit detectable anti-OVA IgG1 titers until after the 2^nd^ administration, a single administration of 100- or 200-cycle ALTA^®^ resuspended in liquid OVA-Alhydrogel at a 1:1 ratio accelerated anti-OVA IgG1 response kinetics and a significantly improved the integrated anti-OVA IgG1 response over time (Figure 4A-4B). While antibody production is certainly a critical response to vaccination, cellular immune responses may also provide protection against many pathogens and were thus of interest to measure. At the 200 ng OVA dose tested, vaccination with Alhydrogel-adjuvanted liquid OVA using a traditional two-dose prime and boost schedule did not generate an antigen-specific CD8+ T cell response (Figure 4C). Incorporating 100- or 200-cycle ALTA^®^ OVA material into the dose, in lieu of a second liquid administration, elicited a more robust antigen specific CD8+ T cell response than two administrations of the Alhydrogel-adjuvanted liquid (Figure 4C). The single administration approach containing both liquid and ALTA^®^ coated particles resulted in circulating OVA-specific CD8+ T cells detectable at nearly 6 months post-injection (Figure 4C). These data support the benefit of one potential scenario for achieving a single administration product with ALTA^®^ by mixing an ALD-coated product with a liquid vaccine formulation with the intent to mimic a traditional, two-dose prime/boost vaccination schedule.

**Figure 4.**
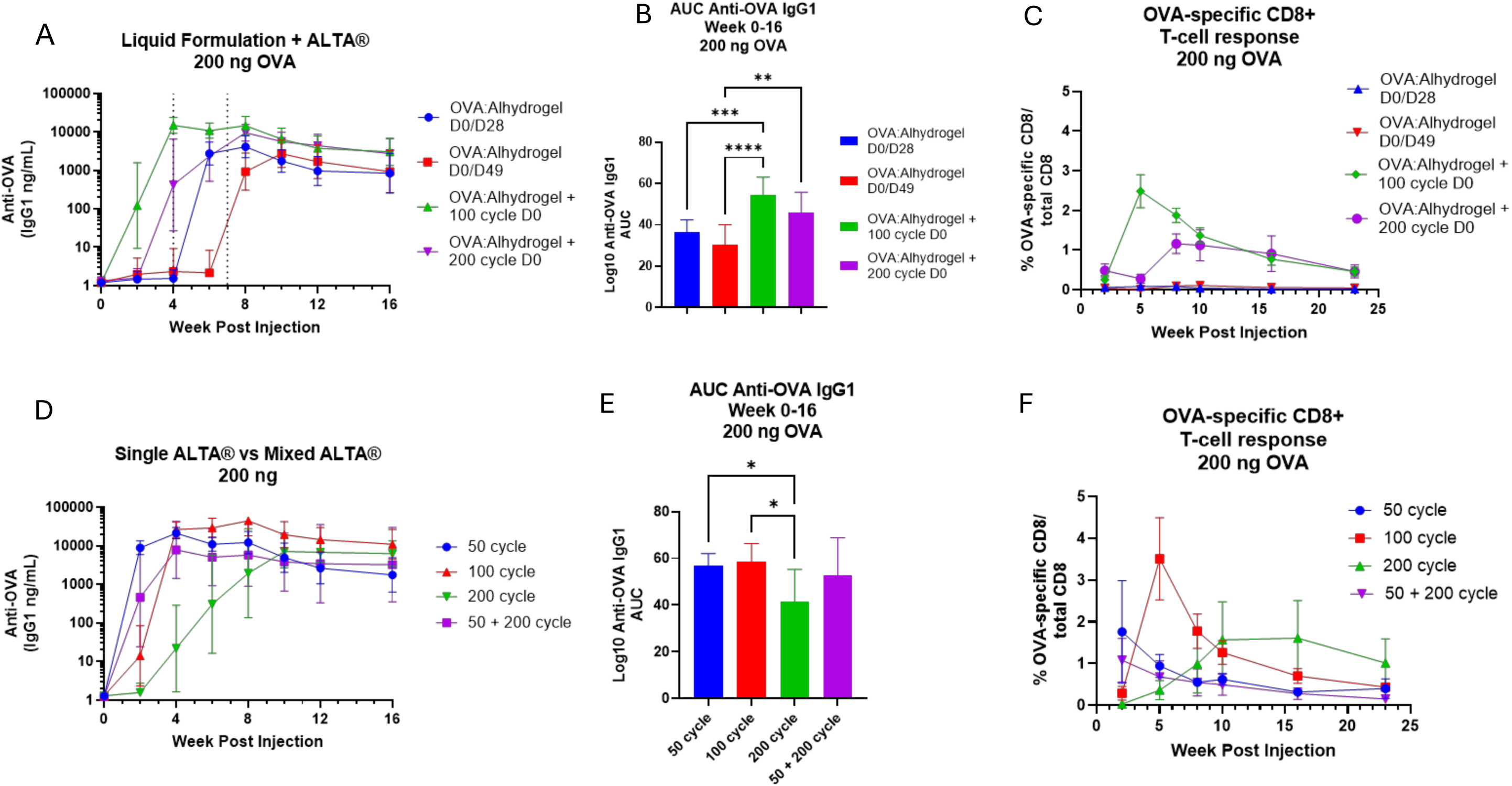
– Comparing immune response from liquid prime/boost paradigm to single ALTA^®^ or mixed ALTA^®^ vaccine products. A. Anti-OVA IgG1 titers following administration of 200 ng OVA dose given from Alhydrogel-adsorbed OVA liquid prime/boost (injection schedule 100 ng D0/100 ng D28 or 100 ng D0/100 ng D49). For mixed products (green/purple traces) 100- or 200-cycle ALTA^®^ powders containing a 100 ng OVA dose were resuspended in a diluent containing 100 ng Alhydrogel-adsorbed OVA and given as a single injection on D0. Plots shows geometric mean titer (n=8-10/group) with 95% confidence interval. B. Total AUC of log10-transformed anti-OVA IgG1 titers shown in A from week 0 to week 16. Column shows mean AUC +/- SEM (n=8-10 mice/group) Data analyzed using one-way ANOVA with Tukey’s multiple comparisons test. **** = p < 0.0001. *** = p < 0.001. ** = p < 0.01. C. The percentage of OVA-specific CD8+ T cells relative to total CD8+ T cells in whole blood was measured using flow cytometry (see methods). Plot shows mean +/- SEM (n=5 mice/group). D. Anti-OVA IgG1 titers following a 200 ng OVA dose given in 50-, 100- or 200-cycle ALD coated vaccine powder. For the mixed ALD vaccine product (purple), 50-cycle and 200-cycle materials were resuspended in diluent at a 2x concentration, then mixed at a 1:1 ratio immediately prior to injection to deliver a total dose of 200 ng OVA. Plot shows geometric mean titer (n=9-10/group) with 95% confidence interval. E. Total area under the curve of anti-OVA IgG1 titer was calculated for all animals after vaccination with 200 ng OVA from 50-, 100-, 200- or mixed 50+200-cycle ALTA^®^ OVA powders. Plot shows the group mean AUC +/- SEM (n=9-10/group). Data analyzed using one-way ANOVA with Tukey’s multiple comparisons test. * = p < 0.05. F. The percentage of OVA-specific CD8+ T cells relative to total CD8+ T cells in whole blood was measured using flow cytometry (see methods). Plot shows mean +/- SEM (n=5 mice/group).

Next, we wanted to evaluate whether the sustained antigen release from ALTA^®^ products alone could elicit a similarly robust immune response. As expected, at a specific antigen dose (200 ng OVA) the kinetics of anti-OVA IgG1 responses were delayed by increasing ALD-cycle number, with 50-cycle material eliciting peak IgG1 titers at week 2 post-administration, while 100-cycle and 200-cycle ALTA^®^ material elicited peak IgG1 responses at week 4, and week 10 post-administration, respectively (Figure 4D). Due to the delayed humoral response kinetics observed, the integrated anti-OVA IgG1 response to 200-cycle ALTA^®^ was significantly lower than 50-cycle and 100-cycle material over the 16 week duration of this study (Figure 4E). The 50-, 100- and 200-cycle ALTA^®^ materials alone, with no liquid vaccine present in the dose, generated robust anti-OVA CD8+ T cell responses with a similar magnitude to what was observed when 100- and 200-cycle ALTA^®^ products were resuspended in a liquid vaccine formulation (Figure 4C + 4F). The kinetics of the anti-OVA CD8+ T cell response elicited by single ALTA^®^ products followed a similar trend as the anti-OVA IgG1 titers, with time to peak anti-OVA CD8+ response delayed by increased ALD-coat thickness, consistent with the antigen release kinetics described previously (Figure 4F, Figures 1-2).

When 50-cycle and 200-cycle ALTA^®^ OVA was mixed at a 1:1 ratio, we observed no difference in anti-OVA IgG1 response compared to 50- or 100-cycle powder given as a standalone product (Figure 4D-4E). However, the mixed 50- and 200-cycle material did elicit a significantly greater integrated anti-OVA IgG1 response than 200-cycle material alone, likely due to the presence of 50-cycle ALTA^®^ in the dose (Figure 4E). The anti-OVA CD8+ T cell response elicited by mixed 50- and 200-cycle ALTA^®^ material exhibited similar magnitude and kinetics as 50-cycle ALTA^®^ alone (Figure 4F).

To provide further insight into the performance of mixed 50- and 200-cycle ALTA^®^ products, we quantified antigen release from 50- and 200-cycle ALTA^®^ containing fluorescently labeled OVA mixed at a 1:1 ratio. In this scenario, we expect the 50-cycle and 200-cycle materials to each contribute 50% of the total observed fluorescent signal. Similar to 50-, 100-and 200-cycle ALTA^®^ products alone (Figure 1A-1B), we observed dose-dependent kinetics of signal loss from mixed 50- and 200-cycle ALTA^®^ particles, with the fluorescent signal reaching 0% at the site of injection on D94 at 2 mg/mL, and D78 at 0.4 mg/mL (Supplementary Figure 6). We observed 50% signal loss by D20 and D14 post injection, at 2 mg/mL and 0.4 mg/mL doses, respectively, which is consistent with time to complete release for 50-cycle material alone (Figure 1A-1B, Supplementary Figure 6). Subsequently, we observed a lower, sustained rate of signal loss, similar to the kinetics observed from 200-cycle ALTA^®^, suggesting that when mixed, 50-cycle and 200-cycle materials behave independently, and the underlying release rates of each product remain unchanged. Given the observed sustained antigen delivery, without temporal delay, from mixed 50- and 200-cycle material, it may be challenging to achieve bi-phasic antigen delivery that results in an anamnestic IgG1 response using mixed ALD-coated vaccine products. However, given the robust immune response elicited by standalone and mixed ALTA^®^ materials (Figure 4D-4F, Supplementary Figure 6), the unique, sustained antigen delivery from ALTA^®^ products may obviate the need for a temporally delayed antigen boost. Together, our results clearly demonstrate that single administration ALTA^®^ vaccine products have the potential to elicit significantly improved humoral and cellular responses relative to two-dose, prime/boost vaccination with a liquid formulation.

### Using ALTA® technology to create a thermostable, extended-release vaccine containing the germline-targeting HIV-1 antigen, N332-GT5

The unique, sustained antigen delivery profile from ALD-coated vaccines may hold benefits against difficult to target pathogens such as HIV-1, where sustained antigen delivery has been shown to improve the immune response to vaccination[11–13]. Recently, it was reported that slow-release delivery of the recombinant HIV-1 Envelope-based germline-targeting antigen, N332-GT5 gp140, improved antigen retention in germinal centers, resulting in an improved overall humoral response[15]. A liquid formulation of N332-GT5 gp140, adjuvanted with SMNP, is currently being tested in an ongoing clinical trial where a fractionated, escalating-dose prime approach is being used in comparison with traditional two-dose, bolus administration to evaluate whether extending antigen delivery via multiple administrations improves the clinical response to vaccination[6]. Given the desire to provide prolonged delivery of N332-GT5 gp140, this antigen may be well-suited to the extended-release profile conferred by ALTA^®^ formulation. Furthermore, designing the vaccine dose(s) within an ALTA^®^ product may have benefits from a manufacturing and stability standpoint as our spray-dried technology platform may impart improved antigen stability and reduce cold-chain storage requirements compared to liquid formulations. Therefore, we manufactured 50-cycle ALTA^®^ products containing unlabeled or fluorescently labeled N332-GT5 gp140 to evaluate the humoral response elicited by ALTA^®^ N332-GT5 gp140 and evaluate antigen release kinetics using a clinically relevant protein subunit antigen rather than a model antigen such as OVA.

Prior to initiating *in vivo* studies, we first confirmed the stability of N332-GT5 gp140 in 50-cycle ALTA^®^ material, as this antigen may be more unstable through our production processes than the model antigen OVA. We observed that 50-cycle ALTA^®^ N332-GT5 gp140 and process intermediates maintained conformational stability throughout the manufacturing process when evaluated with an ELISA utilizing a bnAb against HIV-1, BG18[16] (Figure 5A). Stability of 50-cycle ALTA^®^ N332-GT5 gp140 was also maintained at refrigerated conditions (2-8°C), controlled room temperature (25 ± 2°C/60 ± 5% RH) and accelerated (40 ± 2°C/75 ± 5% RH) stability conditions for up to three months based on ELISA potency data (Figure 5B). These data demonstrate that ALTA^®^ formulation can impart thermostability upon physiologically relevant antigen payloads that may be unstable in traditional liquid formulations.

**Figure 5.**
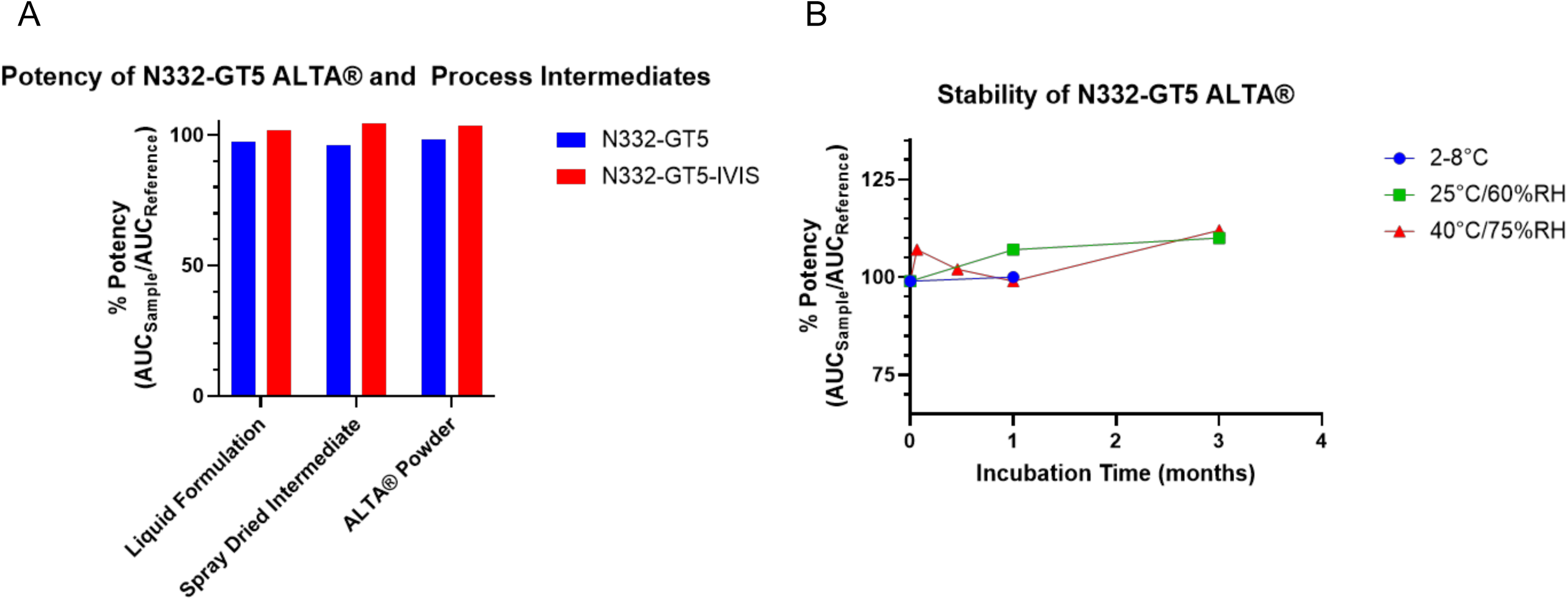
– Conformation of unlabeled and fluorescently labeled N332-GT5 gp140 is maintained through ALTA^®^ manufacturing process and at accelerated stability conditions up to three months. A. Analysis of unlabeled N332-GT5 gp140 or fluorescently labeled N332-GT5 gp140 (N332-GT5-IVIS) potency. B. Potency of unlabeled N332-GT5 gp140 is maintained after 3 months of incubation at conditions up to 40°C/75%RH.

Following vaccination with 50-cycle ALTA^®^ containing unlabeled N332-GT5 gp140, we observed an anti-N332-GT5 gp140 total IgG response at all doses tested, with increased serum antibody titers elicited at higher antigen doses (Figure 6A). Using longitudinal *in vivo* imaging, we found that antigen release from fluorescently labeled 50-cycle ALTA^®^ N332-GT5 gp140 occurred for 28-42 days post-administration, depending on the dose administered, with similar immune response kinetics as the unlabeled ALTA^®^ N332-GT5 gp140, suggesting that the labeled and un-labeled ALTA^®^ products release antigen in a similar manner (Figure 6B). The kinetics of signal loss were consistent with what we observed using 50-cycle ALTA^®^ OVA, confirming the extended antigen release profile of ALD-coated vaccine powders applies across multiple antigen payloads (Figure 1A-1B, Figure 6B).

**Figure 6.**
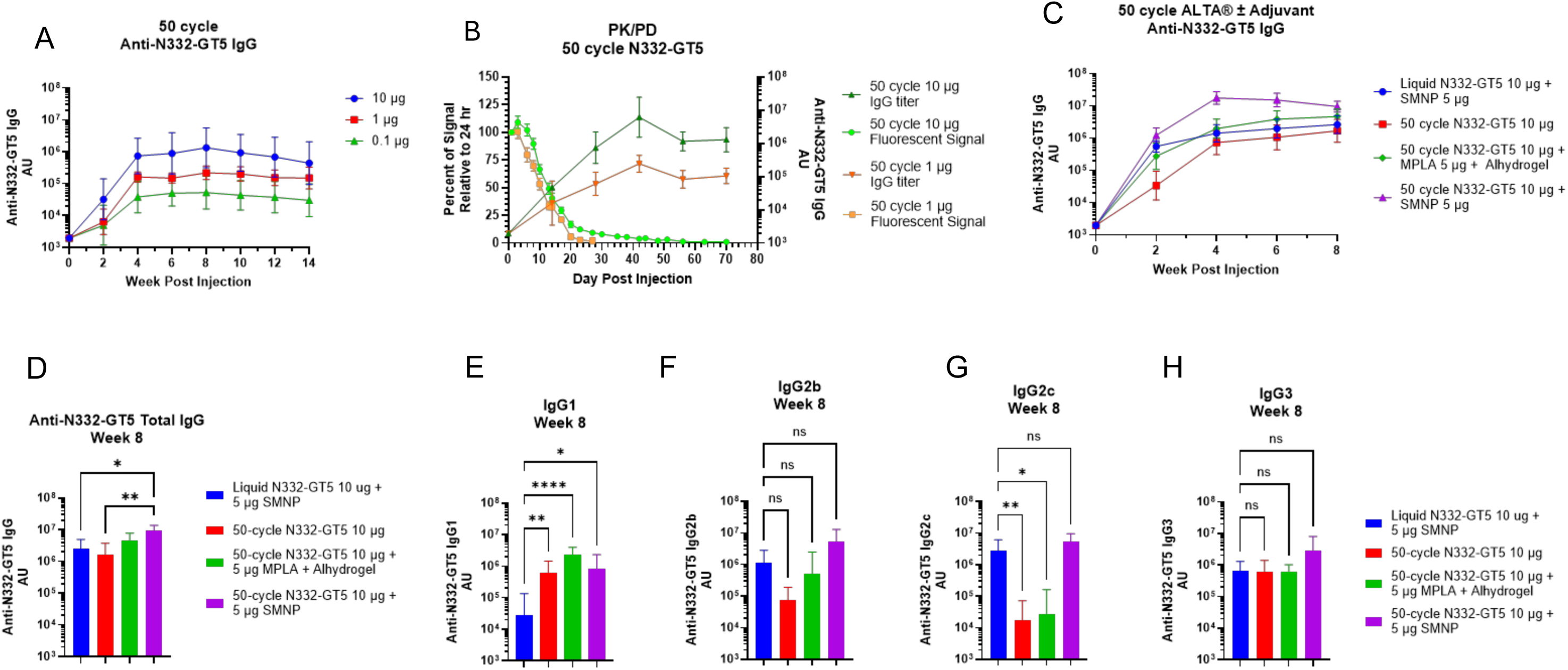
– ALTA^®^ extended-release vaccine containing the germline-targeting HIV-1 immunogen N332-GT5 gp140 outperforms a single administration of the liquid vaccine formulation. A. Total anti-N332-GT5 gp140 IgG titers after a single administration of 50-cycle ALTA^®^ on D0 at indicated doses of N332-GT5 gp140. Plots shows geometric mean titer (n=8/group) +/- 95% confidence interval. B. Analysis of fluorescent signal at site of injection following administration of 50-cycle ALTA^®^ containing fluorescently-labeled N332-GT5 gp140. Plot shows mean +/- SEM of percent of fluorescent radiant efficiency (p/s)/(µW/cm^2^) relative to radiant efficiency at 24 hr timepoint measured at site of injection for 50-cycle ALTA^®^ N332-GT5 gp140 administered at specified doses (square traces). Total anti-N332-GT5 gp140 IgG titers plotted as geometric mean +/- 95% confidence interval (n=5 mice/group) (triangle traces). C. Kinetics of total anti-N332-GT5 gp140 IgG titers elicited by a single administration of 10 µg N332-GT5, delivered in a liquid formulation (blue) or in 50-cycle ALTA^®^ products with/without the presence of adjuvants in the injection diluent. Plot shows geometric mean titer +/- 95% confidence interval (n=16 mice/group red trace, n=8 mice/group all others) D. Total anti-N332-GT5 gp140 IgG1, IgG2b, IgG2c and IgG3 titers elicited by a single administration of 10 µg N332-GT5 gp140 at week 8 post injection, delivered in a liquid formulation (blue) or in 50-cycle ALTA^®^ products with/without the presence of adjuvants in the injection diluent. Plot shows geometric mean titer +/- 95% confidence interval (n=16 mice/group red trace, n=8 mice/group all others). ** = p < 0.01, * = p < 0.05, data analyzed using Kruskal-Wallis test. E-H. Total anti-N332-GT5 gp140 (E) IgG1, (F) IgG2b, (G) IgG2c and (H) IgG3 titers elicited by a single administration of 10 µg N332-GT5 gp140 at week 8 post injection, delivered in a liquid formulation (blue) or in 50-cycle ALTA^®^ products with/without the presence of adjuvants in the injection diluent. Plot shows geometric mean titer +/- 95% confidence interval (n=16 mice/group red trace, n=8 mice/group all others. **** = p < 0.0001, ** = p < 0.01, * = p < 0.05, ns = p > 0.05, data analyzed using Kruskal-Wallis test.

While those initial studies were performed without the addition of adjuvant to ALTA^®^ material to demonstrate our capacity to manufacture immunogenic 50-cycle ALTA^®^ N332-GT5 gp140, it is well established that incorporation of an adjuvant into vaccine formulation can improve the humoral response. Next, we compared the antibody response to a single administration of 10 µg N332-GT5 gp140 + 5 µg SMNP in a liquid formulation to the response elicited by a 10 µg dose of ALTA^®^ N332-GT5 gp140, where SMNP or Alhydrogel-adsorbed MPLA (an AS04-like adjuvant) was administered in the diluent for ALTA^®^ N332-GT5 gp140 injection. We found that while unadjuvanted and MPLA-adjuvanted ALTA^®^ N332-GT5 gp140 elicited a total IgG response with similar magnitude and kinetics as the liquid N332-GT5 formulation, incorporating SMNP to the diluent of ALTA^®^ N332-GT5 gp140 improved the antigen-specific total IgG response, resulting in significantly higher anti-N332-GT5 gp140 serum IgG titers 8 weeks post vaccination (Figure 6C-6D). In previous studies, SMNP has been shown to promote a broad antibody response, with higher levels of antigen specific IgG1 and IgG2a reported relative to various other adjuvants tested[8]. Here, we found that ALTA^®^ N332-GT5 gp140, with or without additional adjuvant, elicited significantly higher anti-N332-GT5 gp140 IgG1 titers than the liquid formulation (Figure 6E). We also found that there was not a significant difference in the magnitude of antigen specific IgG2b and IgG3 responses across all treatment groups (Figure 6F-6H). While the presence of SMNP in the liquid vaccine drove a higher IgG2c response compared to un-adjuvanted and MPLA-adjuvanted ALTA^®^ N332-GT5 gp140, there was no difference in antigen-specific IgG2c levels when ALTA^®^ N332-GT5 gp140 was administered with SMNP (Figure 6G). Thus, incorporation of an additional adjuvant into the ALTA^®^ vaccine dose at the time of administration appears to improve the overall humoral response. Together, these data demonstrate that ALTA^®^ technology may be leveraged to create extended-release products that elicit improved humoral responses compared to liquid vaccine formulations for difficult to vaccinate targets like HIV-1.

## Discussion

There is significant interest in utilizing sustained antigen delivery as an approach to improve immune response to vaccination. Our results shed light on the ability of ALD-coated particle vaccines to elicit strong antigen-specific immune responses, as we found that ALD-coated particles continuously released antigen over weeks/months with a variable release rate. Interestingly, small molecules coated with metal oxides using ALD processes also exhibit pharmacokinetics that are suggestive of sustained, variable rate release[17]. The observation of sustained release featuring a noticeable shift in release rate, that can be delayed by increasing ALD-coat thickness, was consistent across two independent *in vivo* imaging modalities using powders coated with a range of ALD-cycle numbers, suggesting that this is a genuine feature of ALD-coated vaccines. Furthermore, our results contextualize the observed delayed immune responses to higher ALD-cycle number vaccines, as we found that both the early release rate and peak release rates are reduced by increasing alumina-coat thickness. This is suggestive of a model where the timing of the immune response is a function of (daily percent release) x (total particle number/antigen dose administered). At low antigen doses (i.e. low particle numbers), we observe delayed seroconversion, possibly because a higher percentage of particles must release antigen to trigger an immune response. Conversely, at high antigen doses (high particle numbers), we observe earlier seroconversion, as antigen release from a small percentage of the total number of particles administered may provide enough antigen to trigger the earlier onset of immune response.

Notably, the early release rate was non-zero in all studies assessing particle release, regardless of ALD-coat thickness. We speculate that this may be a basic biological feature, as in separate studies we have demonstrated that injection of ALTA^®^ material triggers inflammatory cell recruitment and antigen uptake, potentially through particle phagocytosis, at the site of injection (Ivanova D.I. et al., manuscript in prep). Once engulfed in the phagosome, an individual alumina-coated particle is likely to be exposed to a highly degradative environment that could potentially accelerate ALD-coat rupture and antigen release. Additionally, there is likely a second mechanism of particle release occurring, independent of cellular uptake, as physical or chemically induced degradation of the alumina shell may eventually lead to particle rupture resulting in extracellular antigen release. Naturally, there will be some variation in both spray-dried particle size and ALD-coat quality, even for highly uniform, well-coated ALTA^®^ materials. This variation in particle size and coat quality likely provides a population of particles that readily undergo shell rupture following *in vivo* administration, even for materials coated with 100-200 ALD cycles. It is likely that these two release mechanisms occur in tandem, resulting in particle release immediately following injection.

A consequence of this early release phase is that achieving a temporal delay in antigen release from materials of different ALD-coat thickness may be more challenging than initially expected. While it is possible that high ALD-cycle materials may be delivered at a dose that renders early release negligible, this approach may be limited, as reducing the total dose will result in lower antigen release during the peak release phase. This aspect of sustained particle release may also explain why we observed prolonged maintenance of IgG1 titers when administering doses containing powders of mixed ALD-cycle numbers, rather than an anamnestic response as observed with a traditional prime/boost liquid vaccination (Figure 4D, Supplementary Figure 6).

There are three potential dose delivery paradigms using ALD-coated vaccine powders given as a single shot vaccine: (1) administer a full vaccine dose using material coated with a specific ALD-cycle number, dose-matched to the antigen dose given in a multi-dose liquid prime/boost vaccine, (2) resuspend ALD-coated vaccine powder in a liquid vaccine, thereby administering a liquid priming dose and leveraging the sustained release from ALD-coated material to provide a boost, and (3) administer ALD-coated vaccine products with different ALD-coat thicknesses (i.e. 50-cycle and 200-cycle) with the aim of leveraging the variable release rates to achieve a bi-phasic antigen release and mimic a traditional prime/boost vaccine schedule. A caveat to paradigms 2 and 3 is that the pharmacokinetics of antigen release from ALTA^®^ materials have been shown in these studies to differ from a traditional liquid vaccine formulation. Therefore, it may not be feasible for ALD-coated vaccine products to mimic the pharmacodynamic response elicited by a traditional liquid prime/boost vaccination regimen.

Our results indicate that the sustained antigen delivery from ALTA^®^ vaccines can generate a humoral response that is comparable, or potentially superior to, a traditional multi-dose regimen of a liquid vaccine. Given the unique, sustained antigen release kinetics from ALD-coated vaccines, careful consideration and testing will be required when designing dosing strategies, as it may not be necessary to incorporate a temporally delayed boost dose in ALTA^®^ vaccine products. However, the variable release rates of materials coated with different ALD-cycle numbers may have utility in certain situations. For instance, multiple products of different ALD-coat thicknesses, each containing a different antigen, could be mixed into a single dose, thereby providing a certain amount of antigen evolution over time as antigens will be released at different rates based on their ALD-coat thickness. This may be useful for improving affinity maturation of antibodies against specific epitope(s) or enabling the maturation of germline precursor antibodies to elicit epitope-specific bnAbs for difficult-to-target pathogens such as HIV-1[9].

To illustrate the potential for ALD-coated vaccines in the context of HIV-1, specifically, we demonstrated the capacity to successfully formulate and manufacture an extended-release vaccine containing the clinically relevant, germline-targeting HIV-1 antigen, N332-GT5 gp140, using our ALTA^®^ platform technology. While un-adjuvanted ALTA^®^ N332-GT5 gp140 elicited a robust immune response, we found that the humoral response was improved when a potent adjuvant, SMNP, was incorporated into the diluent. The potential to include adjuvants within the ALTA^®^ particles is also of interest but was beyond the scope of this study. Our findings will likely hold relevance for the development of ALTA^®^ vaccine products, as certain adjuvants may impart additional improvements in immunogenicity to ALTA^®^ materials, while others may not, as we observed when comparing SMNP to an AS04-like adjuvant in our studies. Furthermore, the incorporation of additional adjuvant, mixed with ALTA^®^ material at the time of administration, may affect the required storage conditions or diluent selection for a potential clinical ALTA^®^ vaccine product.

## Conclusion

This study advances our understanding of the pharmacokinetics of the ALTA^®^ platform, which is a critical step toward developing the technology for the application of thermostable, single-shot vaccines. Our studies show that ALTA^®^ vaccine particles deliver antigen in a unique manner that can be described as sustained release with variable release rate. We observe that the duration and timing of particle release can be controlled by adjusting ALD-coat thickness to generate vaccines with the desired antigen delivery characteristics. Overall, our ALD-coated vaccine platform appears to provide an improved immune response compared to a traditional multi-dose liquid vaccine regimen within a single-shot delivery system that can be applied to clinically relevant antigen/adjuvant combinations. Leveraging these understandings and the unique properties of ALTA^®^ vaccines will aide in the design of future preclinical studies, such as a testing in non-human primates.

## Glossary

ALD: Atomic Layer Deposition
ALTA^®^: Atomic Layering and Thermostable Antigen and Adjuvant
AUC: Area under the curve
bnAb: Broadly-neutralizing antibody
HIV-1: Human Immunodeficiency Virus type 1
MPLA: Monophosphoryl Lipid A
OVA: Ovalbumin
SMNP: Saponin/monophosphoryl lipid A nanoparticle

## Acknowledgements

The authors would like to acknowledge and thank the following contributors: Darrell Irvine provided SMNP, BG18 antibody, shared protocols, and reviewed the manuscript. Mariane B. Melo and Agnes Walsh produced the SMNP and assisted in reagent and protocol sharing. Cisloynny Beauchamp-Perez performed the technical transfer and early protocol development of the N332-GT5 gp140 potency ELISA and Evan Norcross assisted with performing and analyzing N332-GT5 gp140 ELISA data. Matthew Lewis assisted with flow cytometry to evaluate T cell responses. James LeCompte, Emma Palm, Grace Beck, Sarah Adzema, Phillip Reyes, and Edis Cehic contributed to manufacturing the ALTA^®^ products used in these studies. The imaging work was performed at the BioFrontiers Institute’s Advanced Light Microscopy Core (RRID: SCR_018302). The Revvity IVIS Lumina III is supported by NIH MIRA (NIGMS): R35GM147455. Biorender software was used to design the graphical abstract.

## Author Contributions

K.A.S. and S.W.B. conceptualized all studies. K.A.S., H.D., and A.M.G. conducted experiments, generated data and interpreted experimental results. I.R.W., E.M.S., A.B.C., and D.L.I. generated data and interpreted experimental results. A.M.R., E.H., Y.H.W., I.A., Y.H., and L.R.A. provided reagents and generated data. K.A.S. prepared the original draft manuscript. K.A.S., S.W.B., A.K.D., A.M.R., and L.R.A. provided review and editing. B.L.S. supervised all studies.

