## Supplemental Material for "Atomic Layering Thermostable Antigen and Adjuvant (ALTA^®^) platform provides unique antigen delivery system through controlled release to improve immune response to vaccination"

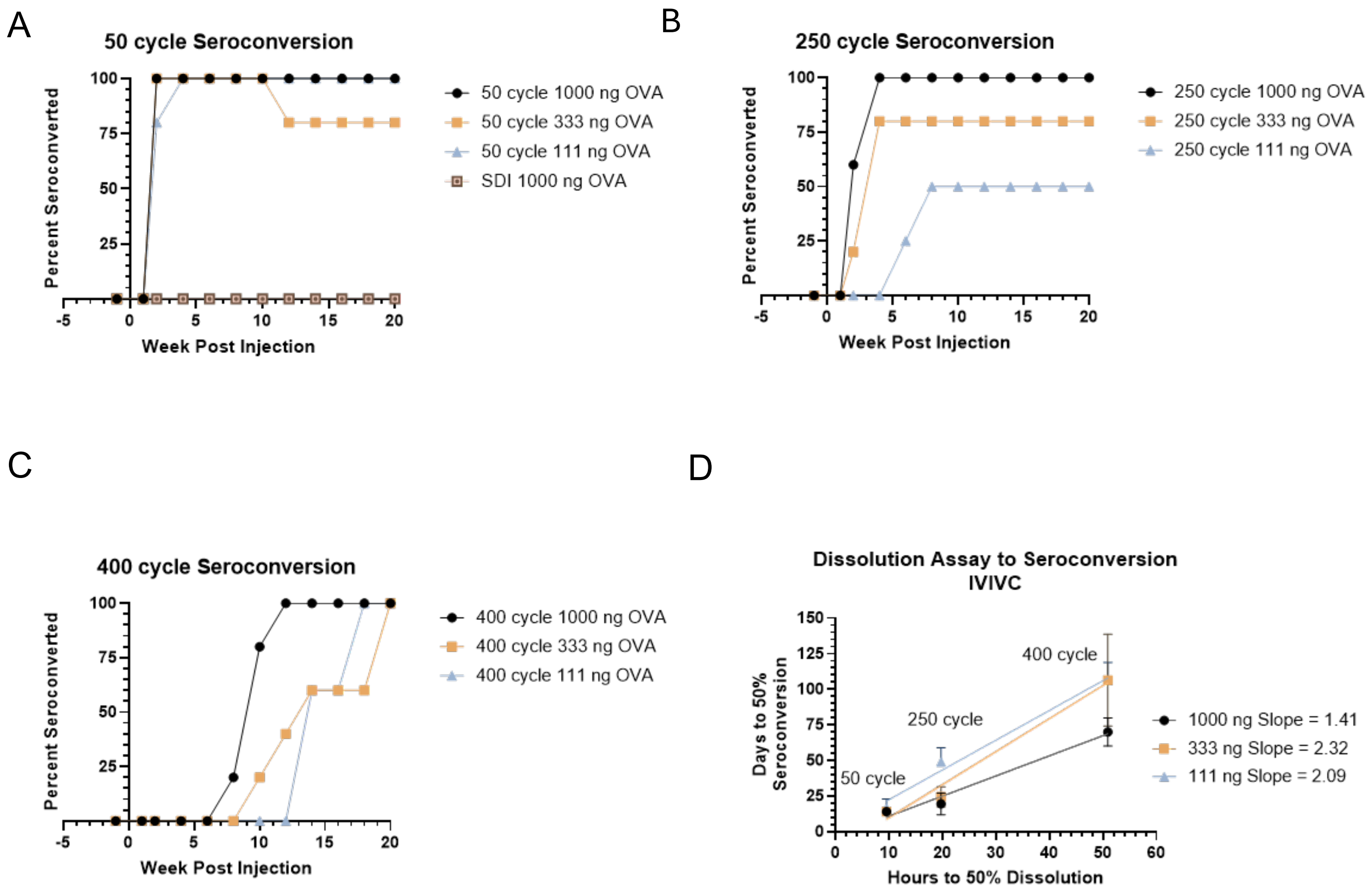


**Supplementary Figure 1 – Delayed antibody responses depend upon coat thickness and dose administered**

A-C. Anti-OVA IgG1 seroconversion percentage, indicating a 2 log-fold increase in anti-OVA IgG1 titers relative to pre-injection baseline, following vaccination with 50-cycle, (B) 250-cycle or (C) 400-cycle ALD coated powders at the indicated OVA doses.

D. Correlation of time to 50% particle dissolution in vitro and days to >50% seroconversion across treated animals in vivo at indicated OVA doses. Simple linear regression of data at each OVA dose shown, with slope of regression line indicated on plot.

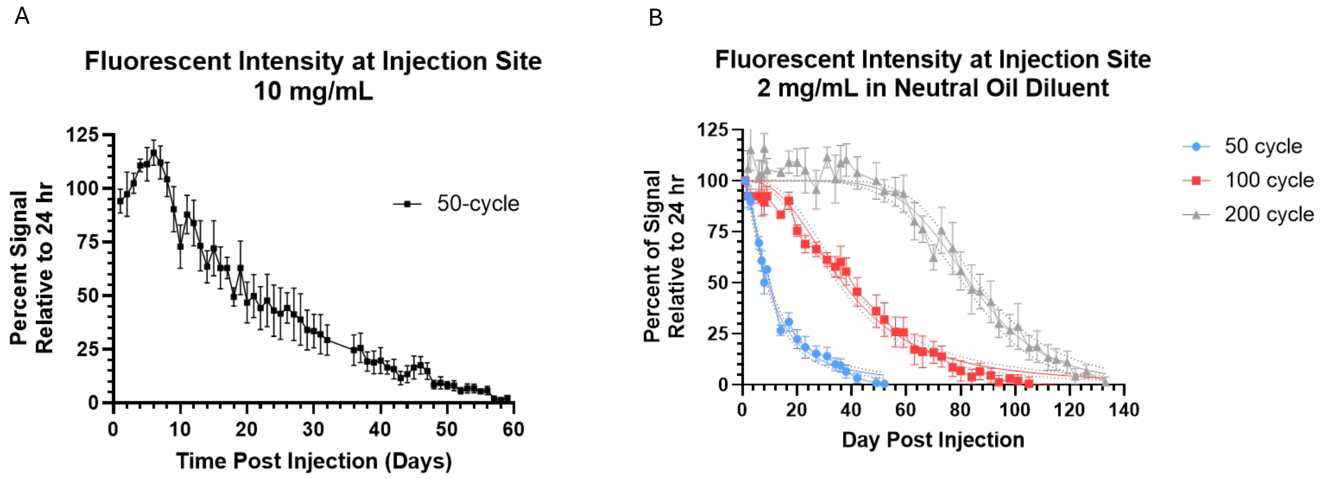


**Supplementary Figure 2 - Increasing dose administered extends ALD-coated particle persistence at injection site**

A. Mean +/- SEM percent of fluorescent radiant efficiency (p/s)/(μW/cm$^2$) relative to radiant efficiency at 24 hr timepoint measured at site of injection for 50-cycle ALD coated powder administered at 10 mg/mL (n=5 mice/group).

B. Mean +/- SEM percent of fluorescent radiant efficiency (p/s)/(μW/cm$^2$) relative to radiant efficiency at 24 hr timepoint measured at site of injection for 50-cycle, 100-cycle and 200-cycle ALD coated powder administered at 2 mg/mL using super refined sesame oil as the diluent for injection (n=5 mice/group).

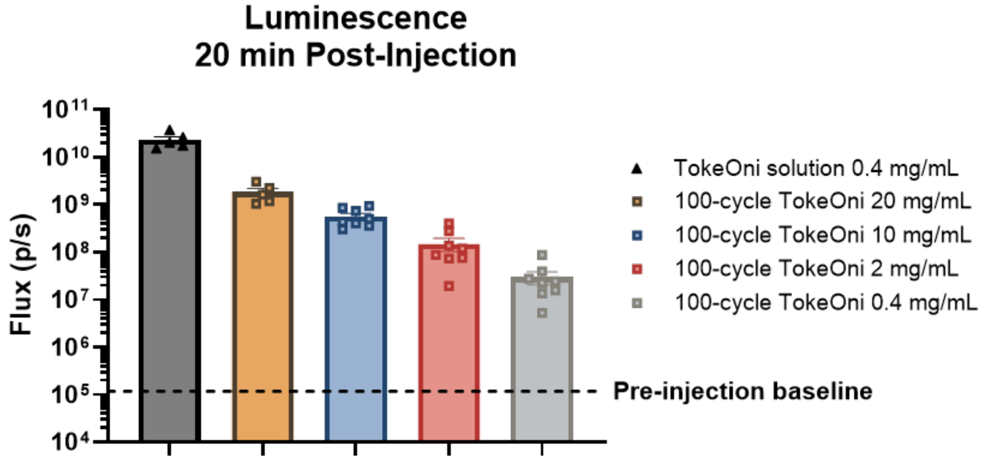


**Supplementary Figure 3 – Activatable probe demonstrates presence of immediately soluble material in ALD coated powders**

Total flux (p/s) measured at the site of injection 20 mins post administration of 100-cycle ALTA® powder containing TokeOni. A 0.4 mg/mL dose of TokeOni solution, matching the total TokeOni dose in 20 mg/mL 100-cycle ALTA® powder, was administered as a control.

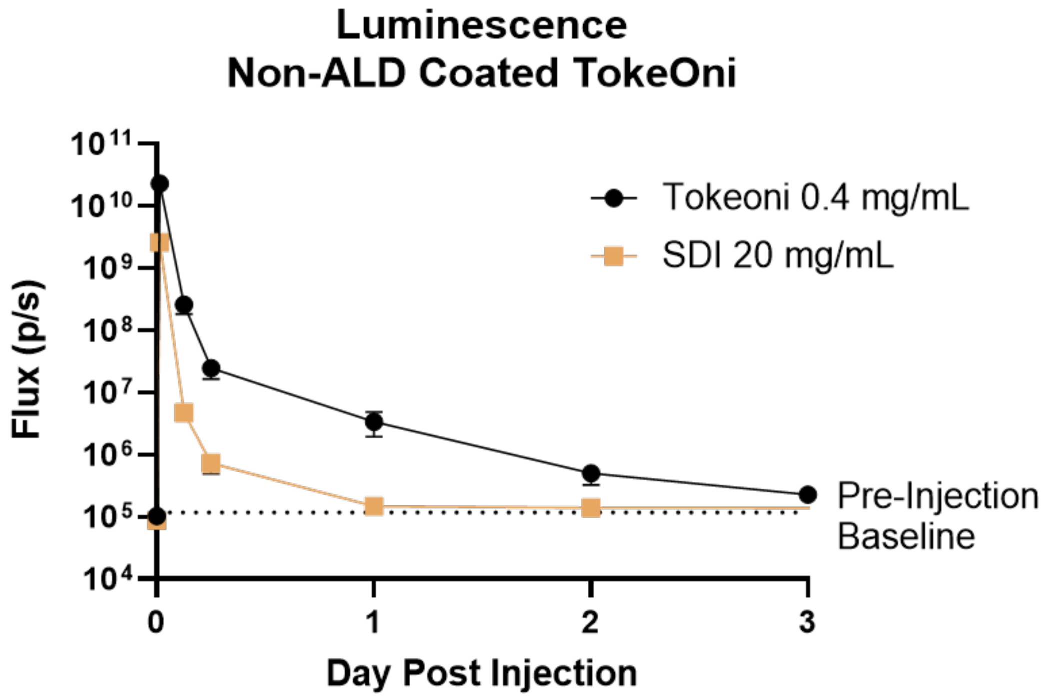


**Supplementary Figure 4 – Duration of luminescent signal from liquid TokeOni solution and uncoated, reconstituted spray dried intermediate (SDI) TokeOni powder**

Total flux (p/s) measured at the site of injection for 3 days following administration of 0.4 mg/mL TokeOni solution or 20 mg/mL uncoated SDI containing TokeOni (pre-cursor to ALD-coated powder) resuspended in sterile saline.

| Sample | Area Size | %Percentage to SDI |
|---|---|---|
| SDI | 53912 | |
| Broken ALTA® | 41478 | 78.7% (corrected for alumina%) |

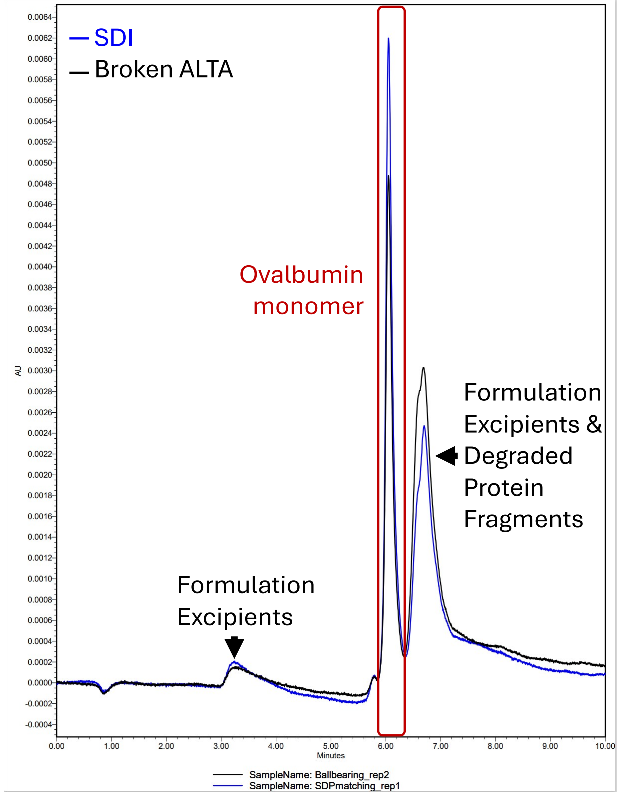


**Supplementary Figure 5 – Uncoated SDI powder and broken 50-cycle ALTA® OVA powder chromatogram showed consistent peak patterns**.

50-cycle ALTA® OVA was physically disrupted, then broken 50-cycle material or uncoated SDI powder was constituted in PBS/0.05% Tween-20 at 10 mg/mL. Size Exclusion Chromatography (SEC) was performed on the Waters ACQUITY UPLC Instrument.

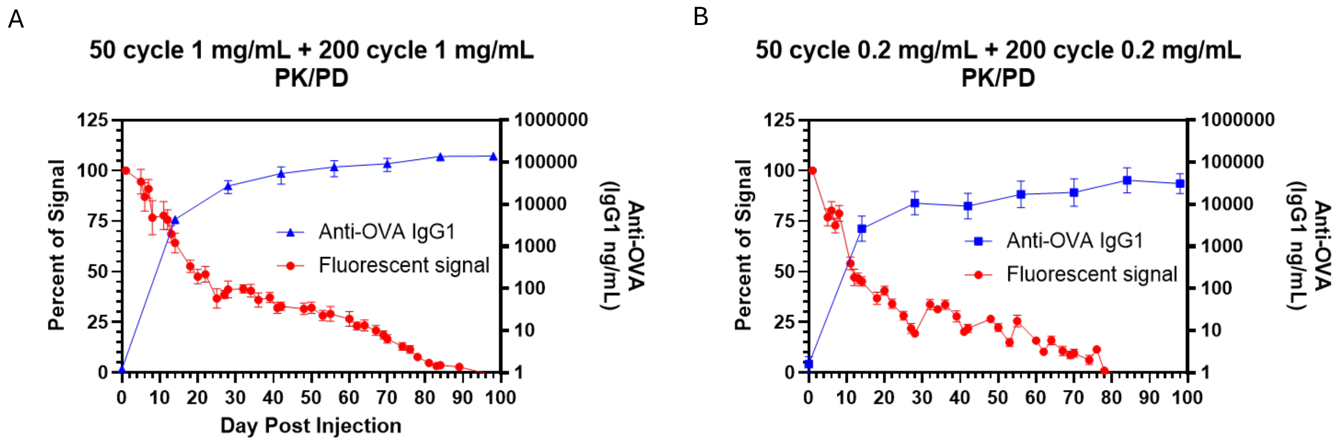


**Supplementary Figure 6 – Mixed ALTA® products demonstrate release kinetics consistent with individual components, and elicit sustained antibody responses**

A-B. Analysis of fluorescent signal at site of injection following administration of 50-cycle and 200-cycle ALTA® containing fluorescently-labeled OVA mixed at a 1:1 ratio dosed at total concentration of 2 mg/mL or (B) 0.4 mg/mL. Plot shows mean +/- SEM of percent of fluorescent radiant efficiency (p/s)/(μW/$cm^2$) relative to radiant efficiency at 24 hr timepoint measured at site of injection (n=5 mice/group) (red traces). Total anti-OVA IgG1 titers plotted as geometric mean +/- 95% confidence interval (n=5 mice/group) (blue traces).

| | 50-cycle | 250-cycle | 400- cycle |
|---|---|---|---|
| A (% Initial Release) | 14% | 2% | 0.4% |
| B (Slope) | 6.2 | 5.1 | 21.0 |
| C (Hrs to Inflection) | 9.5 | 29.2 | 50.9 |
| D (% Full Release) | 94% | 98% | 91% |
| % Alumina | 2.3% | 11.0% | 15.7% |

**Supplementary Table 1 – In vitro dissolution assay results for ALTA® products tested in Supplementary Figure 1**

In vitro dissolution assay 4PL fit parameters and alumina content (% w/w) of ALTA® products coated with 50, 250 or 400 ALD cycles.

| | 50-cycle 2 mg/mL | 50-cycle 0.4 mg/mL | 100-cycle 2 mg/mL | 100-cycle 0.4 mg/mL | 200-cycle 2 mg/mL | 200-cycle 0.4 mg/mL |
|---|---|---|---|---|---|---|
| A - Bottom | 0 | 0 | 0 | 0 | 0 | 0 |
| D - Top | 100 | 100 | 100 | 100 | 100 | 100 |
| C - IC50 (Days to 50% signal loss) | 15.31 | 13.38 | 31.20 | 27.87 | 77.16 | 63.18 |
| B - Hill Slope | -3.226 | -3.261 | -2.408 | -5.062 | -4.893 | -6.076 |
| logIC50 | 1.185 | 1.126 | 1.494 | 1.445 | 1.887 | 1.801 |
| $R^2$ | 0.8704 | 0.8002 | 0.7056 | 0.7899 | 0.6038 | 0.7933 |

**Supplementary Table 2 – 4PL fit parameters of in vivo fluorescent imaging data shown in Figure 1**

4PL analysis constrained to bottom = 0, top = 100. Column headers indicate ALD coat number and dose administered.

| | 50-cycle | 100-cycle | 200-cycle |
|---|---|---|---|
| A (% Initial Release) | 19% | 2% | 0.8% |
| B (Slope) | 2.7 | 9.7 | 16.6 |
| C (Hrs to Inflection) | 10.9 | 15.2 | 24.4 |
| D (% Full Release) | 104% | 99% | 102% |
| % Alumina | 2.3% | 11.0% | 15.7% |

**Supplementary Table 3 – In vitro dissolution assay results for ALTA® products tested in Figure 1**

In vitro dissolution assay 4PL fit parameters and alumina content (% w/w) of ALTA® products coated with 50, 100 or 200 ALD cycles.

| Group | Product 1 | Product 2 | OVA dose Product 1 | OVA dose Product 2 | Administration Schedule |
|---|---|---|---|---|---|
| 1 | 50-cycle ALTA® | | 200 ng | | D0 |
| 2 | 100-cycle ALTA® | | 200 ng | | D0 |
| 3 | 200-cycle ALTA® | | 200 ng | | D0 |
| 4 | 50-cycle ALTA® | 200-cycle ALTA® | 100 ng | 100 ng | D0 |
| 5 | OVA-Alhydrogel 1:50 (D0) | OVA-Alhydrogel 1:50 (D28) | 100 ng | 100 ng | D0/D28 |
| 6 | OVA-Alhydrogel 1:50 (D0) | OVA-Alhydrogel 1:50 (D49) | 100 ng | 100 ng | D0/D49 |
| 7 | OVA-Alhydrogel 1:50 | 100-cycle ALTA® | 100 ng | 100 ng | D0 |
| 8 | OVA-Alhydrogel 1:50 | 200-cycle ALTA® | 100 ng | 100 ng | D0 |

**Supplementary Table 4 – In vivo experimental design related to Figure 4, comparing single administration of ALTA® OVA with two-dose liquid OVA-Alhydrogel formulations**

Table indicates the products administered and the OVA dose contained within each product. Group 5 and Group 6 were given two separate injections of liquid OVA-Alhydrogel, with the second administration occurring on D28 or D49 post-prime, respectively. All other groups received a single administration on D0.
